# SKIPHOS: non-kinase specific phosphorylation site prediction with random forests and amino acid skip-gram embeddings

**DOI:** 10.1101/793794

**Authors:** Thanh Hai Dang, Quang Thinh Trac, Huy Kinh Phan, Manh Cuong Nguyen, Quynh Trang Pham Thi

## Abstract

**Motivation:** Phosphorylation, which is catalyzed by kinase proteins, is in the top two most common and widely studied types of known essential post-translation protein modification (PTM). Phosphorylation is known to regulate most cellular processes such as protein synthesis, cell division, signal transduction, cell growth, development and aging. Various phosphorylation site prediction models have been developed, which can be broadly categorized as being kinase-specific or non-kinase specific (general). Unlike the latter, the former requires a large enough number of experimentally known phosphorylation sites annotated with a given kinase for training the model, which is not the case in reality: less than 3% of the phosphorylation sites known to date have been annotated with a responsible kinase. To date, there are a few non-kinase specific phosphorylation site prediction models proposed.

**Results:** This paper proposes SKIPHOS, a non-kinase specific phosphorylation site prediction model based on random forests on top of a continuous distributed representation of amino acids. Experimental results on the benchmark dataset and the independent test set demonstrate that SKIPHOS compares favorably to recent state-of-the-art related methods for three phosphorylation residues. Although being trained on phosphorylation sites in mamals, SKIPHOS can yield predictions for Y residues better than PHOSFER, a recently proposed plants-specific phosphorylation prediction model.

**Availability and Implementation:** SKIPHOS Web Server is freely available for non-commercial use at http://fit.uet.vnu.edu.vn/SKIPHOS or http://112.137.130.46:5000.

**Supplementary information:** Supplementary data are available at *Bioinformatics* online.

## 1 Introduction

Among known essential post-translation protein modification (PTM) types, phosphorylation is of the top two most common and widely studied one (Khoury *et al*., 2011). A protein kinase catalyzes phosphorylation by adding a phosphate group to certain protein substrates on specific residues, including serine (S), threonine (T), and tyrosine (Y). Phosphorylation is known to regulate most cellular processes such as protein synthesis, cell division, signal transduction, cell growth, development and aging (Hunter, 2000). There are approximately at least 30% of all human proteins are likely to be phosphorylated and about 518 protein kinases encoded in the human genome (Manning *et al*., 2002; Ardito *et al*., 2017). The mouse proteome has more than 540 putative protein kinases (Caenepeel *et al*., 2004) while plant genomes encodes more than 1,000 protein kinases (Vlad *et al*., 2008).

An increasing number of phosphorylation sites in various species have been being experimentally validated, collected and compiled into specialized databases, motivating bioinformatics community to develop advanced in silico prediction models as fast, lower-cost and efficient complements. As a result, various phosphorylation site prediction models have been developed over the past years.

Those models can be broadly categorized as being kinase-specific or non-kinase specific (general). The former aims at building computational models that predicts whether a residue is phosphorylated by a given kinase while the latter to predict irrespective to the kinases. The former thus requires that there are a large enough number of experimentally known phosphorylation sites with a given known catalyzing kinase for the model training. This guarantees the resulting trained models to have satisfactory and significantly persistent kinase-specific phosphorylation predictions. However, note the fact that less than 3% of the phosphorylation sites known to date have been annotated with information about responsible kinases (Newman *et al*., 2013). As a consequence, the number of kinases known to phosphorylate a large amount residues is still limited. To give an example, Phospho.ELM version 9.0 (Dinkel *et al*., 2010), the benchmark dataset for most phosphorylation prediction studies to date (Trost and Kusalik, 2011; Dou *et al*., 2014; Ismail *et al*., 2016; Song *et al*., 2017), has only 9 kinases each catalyzes more than 100 phosphorylation sites. The total residues phosphorylated by such 9 are 1,616, out of 42,500. Over the last decade, an increasing number of non-model organisms’ genomes were sequenced thank to the emerging development of the next generation sequencing technologies, leading to more protein kinases and putative phosphorylaton sites being identified. Therefore, the development of novel non-kinase specific phosphorylation site prediction models is of high demand as an essentially initial phase in phosphorylation studies for a widespread of species. (Trost and Kusalik, 2011).

To date, there are a few non-kinase specific phosphorylation site prediction models proposed. Most of them employs advanced machine learning algorithms, such as neural networks in NetPhos (Blom *et al*., 1999), Support Vector Machines in Musite (Gao *et al*., 2010), PPRED (Biswas *et al*., 2010), PhosphoSVM (Dou *et al*., 2014), and random forests in PHOSFER (Trost and Kusalik, 2013) and RFPhos (Ismail *et al*., 2016). Note that most efforts in the development of phosphorylation prediction models are focused on the kinase-specific (Trost and Kusalik, 2011; Song *et al*., 2017). However, kinase-specific models when being re-adapted for non-kinase specific predictions often generates more false-positives (Dou *et al*., 2014). To this end, this paper introduces a non-kinase specific phosphorylation site prediction model based on random forests on top of a continuous distributed representaion of amino acids. Experimental results demonstrate that our model compares favorably to three recent state-of-the-art methods, namely PhosphoSVM (Dou *et al*., 2014), iPhos-PseEn (Qiu *et al*., 2016) and RFPhos (Ismail *et al*., 2016). Our method out-performs PhosphoSVM, RFPhos and iPhos-PseEn in predictions for S, Y and T residues in terms of overall scoring metrics.

## 2 Materials and methods

### 2.1 Datasets

Experimentally validated phosphorylation sites were extracted from Phospho.ELM version 9.0 (Dinkel *et al*., 2010), the benchmark dataset for most phosphorylation prediction studies to date (Trost and Kusalik, 2011; Dou *et al*., 2014; Ismail *et al*., 2016; Song *et al*., 2017). All redundant protein sequences were eliminated by CD-HIT (Fu *et al*., 2012) with a cutoff of 70% sequence identity. The total number of protein sequences and phosphorylation sites remained for downstream analyses after the redundancy removal are listed in Table 1. For each potential phosphorylation residues (S, Y and T), surrounding windows of certain sizes centering at such are extracted. Resulting subsequences are taken as input to CD-HIT with a 70% cutoff of identity to keep only non-redundant subsequences. A subsequence that has the verified phosphorylation site in the middle is considered a positive, otherwise a negative. In order to avoid the bias, CD-HIT with a 30% identity cutoff were applied to each of both the positive and negative sets to remove redundant corresponding subsequences. Because the numbers of negative subsequences are much larger than that of positive subsequences for S/Y/T (Dou *et al*., 2014), a subset of negative subsequences was randomly selected such that the ratio of negatives to corresponding positives is 1:1 for each S/Y/T. This ratio has been demonstrated to be optimal for phosphorylation site prediction model (Biswas *et al*., 2010). Our dataset (called P.ELM) for cross-validating SKIPHOS is made of these remained non-redundant positive and negative subsequences.

**Table 1.**
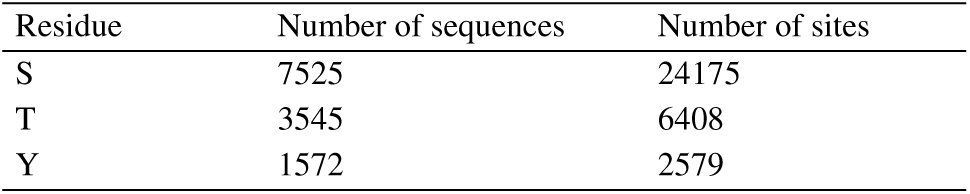
The number of potential phosphorylation sites in non-redundant protein sequences from the benchmark dataset P.ELM

For the independent test set (called PPA), non-redundant *Arabidopsis thaliana* protein sequences from PhosphAt version 3.0 (Zulawski *et al*., 2012) are used to extract experimentally verified positive and negative subsequences. Note that P.ELM contains phosphorylation sites from mammals whereas PPA contains those from *Arabidopsis thaliana*. Such two are shown to be independent of each other (Dou *et al*., 2014). The number of Ser, Thr and Tyr subsequences for specific window sizes in P.ELM and PPA are provided in Table 2. These chosen sizes are the same as in PhosphoSVM and RFPhos, two recent state-of-the-art corresponding methods to which we compared our model.

**Table 2.**
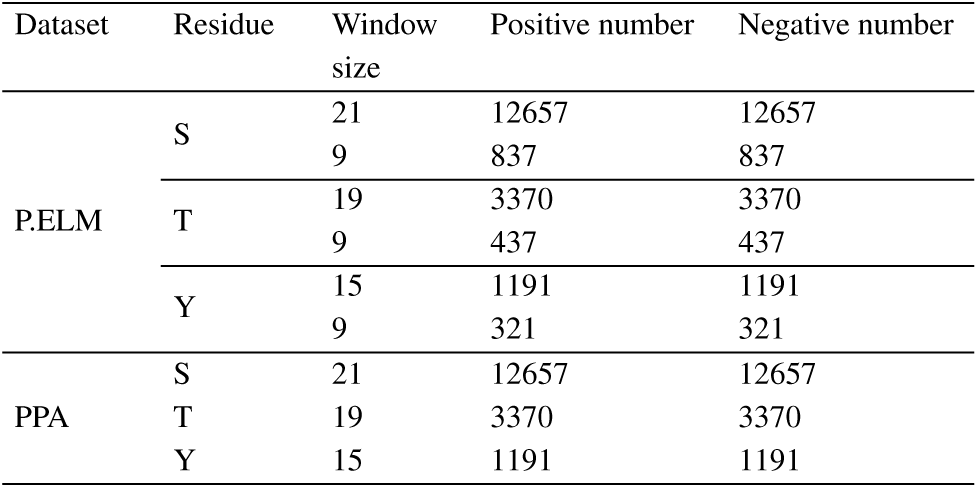
The number of non-redundant known phosphorylation sites for different context window sizes in the benchmark dataset P.ELM and the independent test set PPA

| Dataset | Residue | Window size | Positive number | Negative number |
| --- | --- | --- | --- | --- |
| P.ELM | S | 21 | 12657 | 12657 |
|  |  | 9 | 837 | 837 |
|  | T | 19 | 3370 | 3370 |
|  |  | 9 | 437 | 437 |
|  | Y | 15 | 1191 | 1191 |
|  |  | 9 | 321 | 321 |
| PPA | S | 21 | 12657 | 12657 |
|  | T | 19 | 3370 | 3370 |
|  | Y | 15 | 1191 | 1191 |

### 2.2 Random forests based prediction

Random Forest is a popular ensemble algorithm for classification and regression on high dimensional data (Breiman, 2001). This algorithm constructs a number of decision trees during the training phase and uses the majority vote for prediction. Trees are constructed using bootstrap samples with randomly selected features from the training dataset. The tree construction is guided with the Gini impurity index calculated for each of such selected features. Various recent bioinformatics studies have employed random forests, demonstrating its benefit and robustness for high dimensional datasets (Ismail *et al*., 2016; Song *et al*., 2017).

In this study, random forests with 500 decision trees is used for prediction of phosphorylation sites from rich features derived from subsequences, including: amino acid embeddings, Composition, Transition and Distribution features, Sequence Order Coupling Number features, Quasi Sequence Order features and protein disorder features. The model is implemented using a popular machine learning tool called sklearn (version) (Buitinck *et al*., 2013).

#### 2.2.1 Feature extraction

##### Composition, Transition and Distribution (CTD)

In 1995, Dubchak et al. introduced the Composition, Transition and Distribution (CTD) features for predicting protein folding, which is based on 7 physicochemical properties of amino acids, namely charge, hydrophobicity, normalized van der Waals volume, polarity, polarizability, secondary structure and solvent accessibility (Dubchak *et al*., 1995). Based on a given physicochemical properties, twenty amino acids can be categorized into three groups (i.e. 1, 2 and 3). Each amino acid was then encoded as 1, 2, or 3 according to the group it belongs to. For example, based on the charge property, the subsequence “LLAKKGYQERDLE” is encoded as “1113311123212”. For each such 7 physicochemical properties, there are three types of features could be derived for a subsequence of the length *L* (Li *et al*., 2006; Chou, 2011; Cao *et al*., 2013), including:

- **Composition** of a given group (namely 1, 2 or 3) is the global percentage of such group in the subsequence and is calculated as follows:

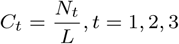

where *N*_*t*_ is the number of times group *t* appears in the subsequence.
- **Transition** for a given pair of groups (*t, v*) characterizes the percent frequency with which group *t* is followed by group *v* or vice versa. It is calculated as follows:

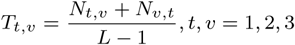

where *N*_*t,v*_ is the number of times group *t* is followed by group *v*.
- **Distribution** descriptor of each group comprises five values, i.e. the fractions of the subsequence where the group is located for the first time, and where 25%, 50%, 75% and 100% of the group are included.

##### Sequence Order Coupling Number (SOCN)

Using the Schneider-Wrede physicochemical distance matrix (Schneider and Wrede, 1994) and chemical distances Grantham matrix (Grantham, 1974), the *k*^*th*^ rank Sequence Order Coupling Number of a *L* amino acid subsequence was calculated as follows:

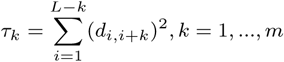

where *d*_*i,i*+*k*_ is the distance between amino acids at position *i* and position *i* + *k, m* = 30 is the maximum lag.

##### Quasi Sequence Order (QSO)

The quasi sequence order comprises two types of features: the first 20 features reflect the frequency ratios of amino acids in a subsequence and the remain reflects the sequence order calculated on the Schneider-Wrede physicochemical distance matrix (Schneider and Wrede, 1994) and the Grantham chemical distance matrix (Grantham, 1974).

The first twenty QSO features are calculated as:

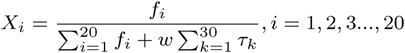

where *f*_*i*_ is the normalized frequency of the amino acid *i, w* = 0.1 is a weighting factor.

The remaining QSO features are given by:

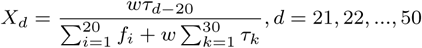

##### Protein Disorder (DIS)

Protein Disorder is an important piece of information of protein functions (Iakoucheva *et al*., 2004). Many phosphorylation prediction studies have used protein disorder as an enriched feature to increase the model accuracy (Gao *et al*., 2010; Ismail *et al*., 2016). In this study, we use DISOPRED (Ward *et al*., 2004) to predict the disorder feature of protein sequences and then the disorder scores predicted for amino acids within a subsequence were extracted.

##### Amino acid embeddings (AAE)

In natural language processing, a word embedding is an algorithm to learn a high-dimensional dense vector representation for words from a very large textual corpus (i.e. training corpus) with billions of words. Words with similar syntax and semantic are embedded to close vectors in the space. It works based on the basic idea that the meaning of a word is affected by surrounding words within its context.

Recently, Mikolov et al. have introduced the Skip-gram model, a novel word embedding architecture based on the neural network language model (Mikolov *et al*., 2013a). Since then, Skip-gram has been employed for numerous natural language processing studies, demonstrating its power and effectiveness in providing good vector representations of words in terms of syntax and semantic relationships.

Given a sentence of N words *w*1, *w*2…, *wN* in the training corpus, the word embedding aims to maximize the probability of observer contexts conditioned on each of such N words at the center:

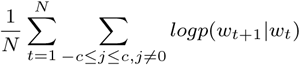

where 2c is the size of the context window centering at *w*_*i*_. The probability *p*(*w*_*t*+*i*_*|w*_*t*_) is calculated by using the softmax function:

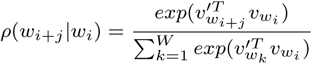

where W is the total number of words in the training corpus,

For the sake of computational efficiency, this full softmax function is approximated with the hierarchical softmax (Morin and Bengio, 2005), in which all W words are represented as leaves of a binary Huffman tree.

The Skip-gram model was then further improved with Negative sampling in which the log probability by the softmax is replaced with the new one as follows (Mikolov *et al*., 2013b):

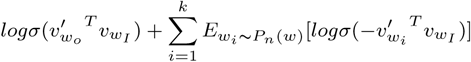

Where the noise distribution *P*_*n*_(*w*) was empirically chosen to be the unigram distribution *U* (*w*) raised to the 3*/*4^*rd*^ power (*i.e*., *U* (*w*)^3*/*4^*/Z*) and *k* is a predefined number of negative samples for each data sample. The authors have experimentally shown that *k* should be in the range 5-20 for small training datasets and 2-5 for large datasets (Mikolov *et al*., 2013b).

In the context of protein bioinformatics, we note that protein sequences or peptides can be considered as “biological” sentences in which each amino acid acts as a distinct “biological” word. Functions of each amino acid on a protein sequence/peptide are affected by neighboring ones surrounding such. In this regard, protein sequences remained after the redundancy removal were used as the training corpus for the Skip-gram. We employ word2vec (Mikolov *et al*., 2013a,b), which implements the state-of-the-art Skip-gram model, to learn continuous vector representations of 300 dimensions for 20 amino acids. This number was experimentally shown to help our proposed model to produce the best performance for S, Y and T phosphorylation predictions.

### 2.3 Model evaluation

10-fold cross validation was implemented for the model evaluation. Five well-known evaluation metrics were used to score the model performance, including the area under the ROC curve (AUC), F1 score, precision, recall and Metthew’s correlation coefficient (MCC), which are given as follows:

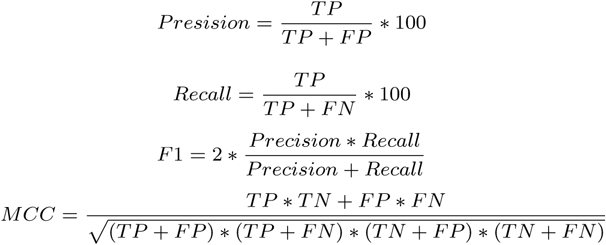

where TP, TN, FP, and FN respectively represent the number of true positives, true negatives, false positives, and false negatives in the fusion matrix. We run a 10-fold cross validation procedure 30 times and the average of resulting aforementioned performance scores are reported for evaluation.

## 3 Results and Discussion

Our proposed general phosphorylation site prediction model SKIPHOS uses subsequences of 9, 15 and 19 amino acids centering at S, Y and T, respectively. Such three lengths have been experimentally demonstrated to produce the best performance correspondingly for S, Y and T when compared with other lengths. We here only present our model’s performance using the length 21, 9 and 9 for S, Y and T respectively as an extra reference for the comparison with two recent state-of-the-art corresponding models, namely RFPhos (Ismail *et al*., 2016) and PhosphoSVM (Dou *et al*., 2014). These extra lengths allow our models to work on subsequences of the same lengths as such two models: RFPhos uses subsequences of 9 amino acids for S, Y and T while PhosphoSVM uses subsequences of 21, 15 and 19 for S, Y and T, respectively. We re-implemented these two models for the (cross-validated) comparison with SKIPHOS on our subsequence datasets. The reasons include: (i) the authors of RFPhos only provide three trained models (for S, Y, and T), coupled with the subsequence dataset on which their models were trained, and (ii) the authors of PhosphoSVM do provide neither and do not release its source code as well.

Experimental results show that our proposed model yields favorable performance on non-kinase specific prediction of S, Y and T phosphorylation sites, when compared to iPhos-PseEn (Qiu *et al*., 2016), RFPhos (Ismail *et al*., 2016) and PhosphoSVM (Dou *et al*., 2014), three recent state-of-the-art corresponding models.

For 10-fold cross validation on the subsequence dataset of RFPhos, SKIPHOS yields excellent prediction performance. It archives the AUC values of 90%, 91.7% and 91.3% for S, Y and T residues, respectively, which are better than those from both RFPhos (i.e. 88%, 91% and 90%) and phosphoSVM (i.e. 84%, 74% and 82%). Futher, when using random forests of 100 decision trees, which is the same number as in RFPhos, SKIPHOS can yield performance with the AUC values of 89.5%, 91.3% and 90.8%, respectively, out-performing RFPhos. This demonstrates the predictive power of SKIPHOS’s features. Note that the RFPhos model implemented by us performs exactly on-par with the trained model given by the RFPhos authors (data not shown), guaranteeing that RFPhos was correctly re-implemented by us.

Further, SKIPHOS is also compared with iPhos-PseEn (Qiu *et al*., 2016), a human-specific non-kinase phosphorylation site predictor based on ensemble random forests. The same 5-fold cross validation scheme as used in iPhos-PseEn is employed for SKIPHOS on the dataset provided by iPhos-PseEn. To this end, SKIPHOS yields excellent performance for S, Y and T in terms of AUCs, i.e. 91.96%, 88.23% and 84.43%, respectively. Prediction results show that, for S and Y, SKIPHOS out-performs iPhos-PseEn and vice versa for T. The prediction accuracy values of iPhos-PseEn are all less than 80% (79.76% for S, 76.28% for Y and 79.88% for T) while those of SKIPHOS are 86.66% for S, 80.52% for Y and 76.28% for T. We note that the MCC values of SKIPHOS are much better than those of iPhos-PseEn (see Table 3 for more details).

**Table 3.** Performance of SKIPHOS and iPhos-PseEn (Qiu *et al*., 2016) on the dataset provided by the iPhos-PseEn’s authors. The best value of each scoring metric is in bold.

| Methods | Residue = Y |  |  |  |  |  |  |
| --- | --- | --- | --- | --- | --- | --- | --- |
|  | F1 | AUC | MCC | Recall | Precision | Accuracy | Specificity |
| SKIPHOS | **0.7939** | **0.8823** | **0.6119** | **0.7680** | **0.8234** | **0.8052** | **0.8353** |
| Iphos-PreEn | 0.7622 | N/A | 0.3244 | 0.7618 | 0.7626 | 0.7628 | 0.7629 |
| Methods | Residue = T |  |  |  |  |  |  |
|  | F1 | AUC | MCC | Recall | Precision | Accuracy | Specificity |
| SKIPHOS | 0.7265 | **0.8443** | **0.5389** | 0.6428 | **0.8383** | 0.7628 | **0.8760** |
| Iphos-PreEn | **0.7322** | N/A | 0.3444 | **0.7151** | 0.7501 | **0.7988** | 0.7618 |
| Methods | Residue = S |  |  |  |  |  |  |
|  | F1 | AUC | MCC | Recall | Precision | Accuracy | Specificity |
| SKIPHOS | **0.8467** | **0.9196** | **0.7024** | **0.8323** | **0.8619** | **0.8510** | **0.8666** |
| Iphos-PreEn | 0.7970 | N/A | 0.3901 | 0.7964 | 0.7975 | 0.7976 | 0.7978 |

For cross-validation on our P.ELM subsequence dataset, SKIPHOS yields good prediction performance for Y (*AUC* = 75.5%) and very good for S (*AUC* = 84.5%) and T (*AUC* = 84.4%). It, however, still out-performs PhosphoSVM (for S, Y and T sites) and RFPhos (for both Y and T) in terms of all aforementioned scoring metrics (see Table 4 for more details).

**Table 4.** Performance of SKIPHOS in comparison with two recent state-of-the-art related models using 10-fold cross-validation on the benchmark dataset P.ELM. (*) indicates the use of the same context window sizes as in SKIPHOS, i.e. 15 for Y and 19 for T. The best value of each scoring metric is in bold.

| Methods | Residue = Y |  |  |  |  |
| --- | --- | --- | --- | --- | --- |
|  | F1 | AUC | MCC | Recall | Precision |
| SKIPHOS | **0.700** | **0.755** | **0.396** | **0.711** | **0.691** |
| RFPhos* | 0.660 | 0.713 | 0.318 | 0.668 | 0.654 |
| PhosphoSVM | 0.627 | 0.677 | 0.253 | 0.628 | 0.627 |
| RFPhos | 0.607 | 0.656 | 0.226 | 0.603 | 0.616 |
| Methods | Residue = T |  |  |  |  |
|  | F1 | AUC | MCC | Recall | Precision |
| SKIPHOS | **0.765** | **0.844** | **0.547** | 0.744 | **0.788** |
| RFPhos* | 0.747 | 0.824 | 0.502 | 0.741 | 0.753 |
| PhosphoSVM | 0.729 | 0.804 | 0.464 | 0.720 | 0.738 |
| RFPhos | 0.747 | 0.815 | 0.475 | **0.784** | 0.716 |
| Methods | Residue = S |  |  |  |  |
|  | F1 | AUC | MCC | Recall | Precision |
| SKIPHOS | 0.765 | **0.845** | 0.521 | 0.785 | 0.749 |
| PhosphoSVM | 0.743 | 0.819 | 0.499 | 0.724 | 0.762 |
| RFPhos | **0.781** | 0.842 | **0.547** | **0.816** | **0.751** |

For S residues, it performs on-par with RFPhos. However, SKIPHOS can predict better than RFPhos at low false positive rate (i.e. <20%) (see Figure 1). It can be argued that the out-performance of SKIPHOS over RFPhos for Y and T may come from the larger context windows used by SKIPHOS. We thus evaluate RFPhos with the larger context windows as used in SKIPHOS, i.e. 15 amino acids for Y and 19 for T. To this end, this variant of RFPhos still performs worse than ours, demonstrating the great utility of features used in SKIPHOS (Table 4).

**Fig. 1.**
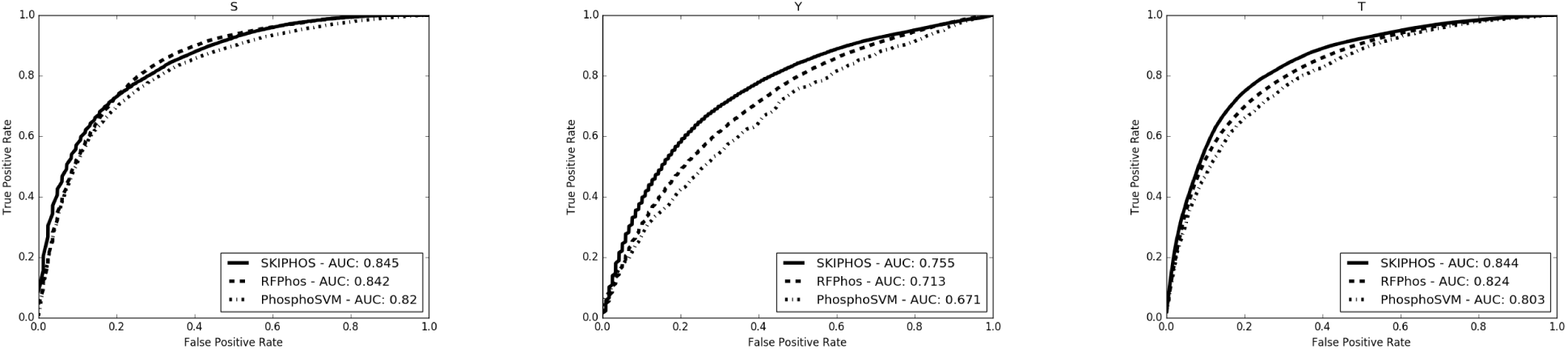
The ROC curves of SKIPHOS and two recent state-of-the-art related models using 10-fold cross-validation on the benchmark dataset P.ELM. (*) indicates the use of the same context window sizes as in SKIPHOS, i.e. 15 for Y and 19 for T.

Interestingly, experimentally results showed that extending the context windows surrounding Y and T brings significant performance improvements to SKIPHOS and RFPhos, as well. It is, however, vice versa for S. This phenomenon suggests that factors determining the phosphorylation status of S residues are likely to be located in the windows of only 9 amino acids centering at them. However, for Y and T residues, these windows are much larger, i.e. 15 and 19, respectively.

We do evaluate the impact of every feature type for SKIPHOS in prediction of non-kinase specific phosphorylation sites by 10-fold cross validating SKIPHOS with such each. Table 5 shows the greatest impact of the amino acid embeddings when they contribute up to 96.6%, 86.5% and 95.3% of the SKIPHOS predictive capacity for S, Y and T, respectively. Among all feature types, the amino acid embeddings contribute most to the predictive strength of SKIPHOS for S and T. For Y, it takes the second place, a little bit after the QUASI. Surprisingly, the contributions of the amino acid embeddings for SKIPHOS in prediction of S and T are on-par with those of all remaining others together. Note that the amino acid embeddings are calculated offline just only for one time while such all remaining features are calculated upon protein sequences. It is useful when using SKIPHOS to make prediction for a newly given protein sequence.

**Table 5.** Performance of SKIPHOS with the use of different feature types. All models are tested with 10-fold cross-validation on the benchmark dataset P.ELM. * indicates that DIS is replaced with QSO in the case of T.

| Feature | Residue=Y |  |  |  |  | Residue = T |  |  |  |  | Residue = S |  |  |  |  |
| --- | --- | --- | --- | --- | --- | --- | --- | --- | --- | --- | --- | --- | --- | --- | --- |
|  | AUC | F1 | Recall | MCC | Accuracy | AUC | F1 | Recall | MCC | Accuracy | AUC | F1 | Recall | MCC | Accuracy |
| AAE | 0.653 | 0.607 | 0.603 | 0.225 | 0.612 | 0.804 | 0.706 | 0.644 | 0.476 | 0.734 | 0.801 | 0.714 | 0.666 | 0.476 | 0.736 |
| AAE+CTD | 0.720 | 0.666 | 0.677 | 0.326 | 0.663 | 0.835 | 0.752 | 0.713 | 0.536 | 0.767 | 0.815 | 0.725 | 0.682 | 0.490 | 0.744 |
| AAE+DIS | 0.677 | 0.632 | 0.620 | 0.281 | 0.640 | 0.803 | 0.718 | 0.677 | 0.475 | 0.736 | 0.809 | 0.727 | 0.690 | 0.489 | 0.744 |
| AAE+QSO | 0.667 | 0.617 | 0.613 | 0.244 | 0.622 | 0.808 | 0.709 | 0.647 | 0.479 | 0.736 | 0.805 | 0.716 | 0.668 | 0.478 | 0.737 |
| AAE+SOCN | 0.661 | 0.613 | 0.608 | 0.239 | 0.619 | 0.807 | 0.707 | 0.643 | 0.478 | 0.735 | 0.804 | 0.716 | 0.669 | 0.478 | 0.737 |
| AAE+CTD+DIS* | **0.748** | **0.694** | **0.707** | **0.382** | **0.690** | **0.838** | **0.754** | 0.714 | **0.541** | **0.769** | **0.824** | 0.741 | 0.711 | **0.509** | **0.754** |
| CTD | 0.565 | 0.580 | 0.587 | 0.157 | 0.578 | 0.581 | 0.582 | 0.581 | 0.171 | 0.585 | 0.659 | 0.637 | 0.651 | 0.264 | 0.632 |
| DIS | 0.616 | 0.588 | 0.587 | 0.182 | 0.591 | 0.649 | 0.631 | 0.666 | 0.228 | 0.613 | 0.645 | 0.651 | 0.729 | 0.233 | 0.612 |
| QSO | 0.676 | 0.630 | 0.639 | 0.256 | 0.627 | 0.769 | 0.715 | 0.723 | 0.430 | 0.715 | 0.766 | 0.712 | 0.730 | 0.417 | 0.708 |
| SOCN | 0.545 | 0.529 | 0.532 | 0.059 | 0.529 | 0.604 | 0.576 | 0.584 | 0.148 | 0.574 | 0.614 | 0.584 | 0.587 | 0.170 | 0.585 |
| All-AAE | 0.739 | 0.683 | 0.685 | 0.368 | 0.684 | 0.801 | 0.741 | **0.758** | 0.476 | 0.737 | 0.810 | **0.751** | **0.787** | 0.484 | 0.741 |

### 3.1 Performance on the independent test set

SKIPHOS, PhosphoSVM re-implemented by us and the trained RFPhos given by its authors are used to make predictions of S, Y and T phosphorylation sites for the independent test set PPA. Predictions by PHOSFER are obtained by submitting all protein sequences in PPA to its freely online available webserver.

For S residues, SKIPHOS achieves the prediction performance with AUC of 69.1% (F1 of 55.2%). They are respectively of 63.4% (62%) and 65.7% (54.3%) for Y and T (see Table 6 for more scoring metrics). The AUC values of RFPhos for S, Y and T are 49.6%, 52% and 52.6%, respectively, demonstrating the outperformance of SKIPHOS over RFPhos. It is also the case for PhosphoSVM when its AUC values for S, Y and T are 66%, 57.8% and 63.1%, respectively.

**Table 6.** Performance of SKIPHOS in comparison with recent state-of-the-art related models on the independent test set PPA. The best values of scoring metrics for the models of the same type (i.e. SKIPHOS, RFPhos and PhosphoSVM) are in bold.

| Methods | Residue= Y |  |  |  |  |
| --- | --- | --- | --- | --- | --- |
|  | F1 | AUC | MCC | Recall | Precision |
| SKIPHOS | 0.620 | **0.634** | **0.197** | 0.658 | **0.586** |
| RFPhos | **0.649** | 0.520 | 0.008 | **0.926** | 0.500 |
| PhosphoSVM | 0.508 | 0.578 | 0.106 | 0.462 | 0.563 |
| PHOSFER | 0.542 | 0.604 | 0.121 | 0.521 | 0.566 |
| Methods | Residue= T |  |  |  |  |
|  | F1 | AUC | MCC | Recall | Precision |
| SKIPHOS | 0.543 | **0.657** | **0.224** | 0.467 | **0.649** |
| RFPhos | **0.667** | 0.526 | 0.000 | **1.000** | 0.500 |
| PhosphoSVM | 0.489 | 0.631 | 0.189 | 0.394 | 0.642 |
| PHOSFER | 0.612 | 0.682 | 0.229 | 0.610 | 0.615 |
| Methods | Residue= S |  |  |  |  |
|  | F1 | AUC | MCC | Recall | Precision |
| SKIPHOS | 0.552 | **0.691** | **0.265** | 0.462 | **0.686** |
| RFPhos | 0.585 | 0.496 | -0.01 | **0.713** | 0.496 |
| PhosphoSVM | **0.624** | 0.660 | 0.237 | 0.632 | 0.615 |
| PHOSFER | 0.714 | 0.807 | 0.390 | 0.773 | 0.663 |

Looking deeper into the ROC curves of 3 models in Figure 2 it can be observed that the ROCs of SKIPHOS for S, Y and T are respectively above those of both RFPhos and PhosphoSVM in the upper left regions, in which recall values are high (let says >= 50%) and false positive rates (FPR) are low (let says <=40%), implying the better performance. Within the lower left regions (recall <=32.5% and FPR <=18%) SKIPHOS performs better than RFPhos and PhosphoSVM, except only for the case of T predicted by PhosphoSVM.

**Fig. 2.**
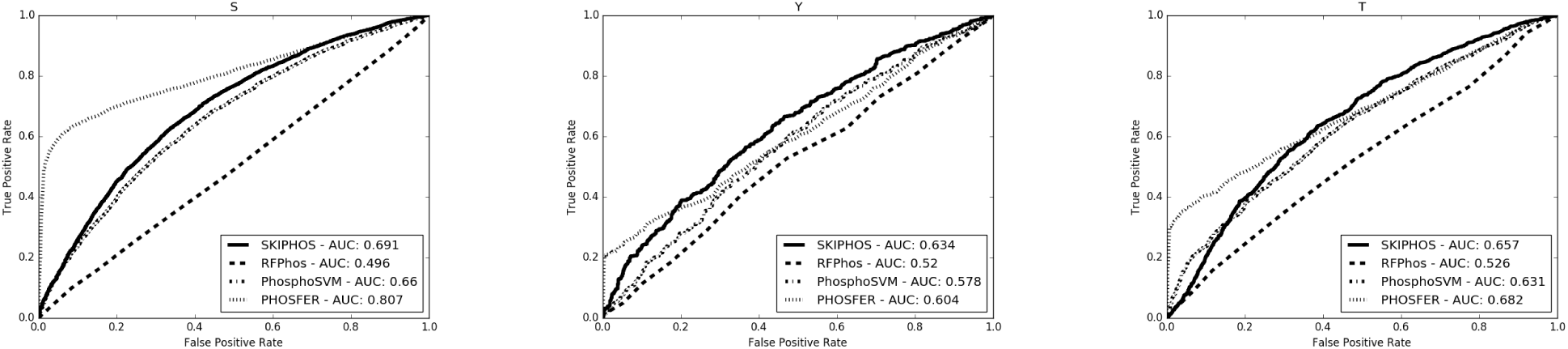
The ROC curves of SKIPHOS and recent state-of-the-art related models on the independent dataset PPA.

Compared with the plant-specific model PHOSFER, the SKIPHOS performance is better only at the recall values greater than 35% (associated with FPR >21%) for Y and only up to 59% (37.5%) for T. The ROC curve of SKIPHOS for S is totally under that of PHOSFER, implying the worse performance, which is not a surprise since PHOSFER are trained from a much larger training dataset from 9 organisms including plants while SKIPHOS is not trained on plant phosphorylation sites. However, this fact, in turn, demonstrates the predictive strength and stability of SKIPHOS when it can yield predictions in the upper left regions (high recall and possibly allowed low FPR) of ROCs that are better than PHOSFER for Y and T (Figure 2).

## 4 Accessibility

We developed a web server with user-friendly graphic interface for SKIPHOS and deployed it online accessible free for non-commercial use at http://fit.uet.vnu.edu.vn/SKIPHOS. Users just only need to provide a protein sequence, choose phosphorylation types for which they want SKIPHOS to make prediction. When completed, the web server returns a list of all phosphorylation residues predicted for each of chosen types.

## 5 Conclusion

In this paper we present SKIPHOS, a novel computational model for non-kinase specific prediction of phosphorylation sites using random forests and amino acid skip-gram embeddings. Experimental results from rigorous validation schemes demonstrate the favorable strength and stability of SKIPHOS when compared to recent state-of-the-art related models, namely PhosphoSVM (Dou *et al*., 2014), iPhos-PseEn (Qiu *et al*., 2016) and RFPhos (Ismail *et al*., 2016). The SKIPHOS performance crossvalidated on the benchmark dataset is better than that of iPhos-PseEn, RFPhos and PhosphoSVM for all cases, except for S residue when being compared with RFPhos, with which on-par performance is observed. However, SKIPHOS outperforms both RFPhos and PhosphoSVM on the independent data set of phosphorylation sites in plants. Surprisingly, SKIPHOS can yield high-recalled predictions for Y and T that are better than those of PHOSHER. Note that PHOSFER is trained on a large dataset containing phosphorylation sites in plants whereas SKIPHOS is only trained on a smaller dataset of those in mammals.

We anticipate that SKIPHOS with a freely available web server will facilitate other basic and/or translational researches related to identification of phosphorylation sites, accelerating discoveries of new important biochemical insights at low costs.

## Acknowledgements

This work has been supported by Vietnam National University, Hanoi (VNU), under Project No. QG.15.21. We would like to thank the author of RFPhos, Dr. Hamid D. Ismail, Research Associate at Department of Animal Sciences, North Carolina Agricultural and Technical State University, for sending us their trained RFPhos model and dataset.

